# Atovaquone-venturicidin A combination is synergistic against *Plasmodium falciparum*

**DOI:** 10.1101/2025.08.18.670846

**Authors:** Dennis Hauser, Sibylle Sax, Sonja Keller, Monica Cal, Marcel Kaiser, Pascal Mäser

**Affiliations:** Swiss Tropical and Public Health Institute, 4123 Allschwil, Switzerland; University of Basel, 4002 Basel, Switzerland

## Abstract

We report the synergistic interaction between the complex III inhibitor atovaquone and an ATP synthase inhibitor, venturicidin A, against *Plasmodium falciparum*. Our results suggest the simultaneous inhibition of the parasite’s primary and alternative pathways for generating the mitochondrial membrane potential to be responsible for this phenomenon, whereby the alternative pathway relies on ATP synthase running in reverse mode. We hypothesize that the synergistic interaction between atovaquone and proguanil could follow a similar mechanism.

---

The prophylactic antimalarial Malarone contains atovaquone and proguanil. In combination, these two drugs have a synergistic effect against *Plasmodium falciparum* (Canfield, 1995). Atovaquone is a complex III inhibitor (Fry, 1992) that affects the mitochondrial membrane potential in *P. falciparum* (Srivastava, 1997; Srivastava, 1999), and indirectly also blocks pyrimidine *de novo* synthesis by preventing the re-oxidation of ubiquinone that is required to make orotate (Painter, 2007). Proguanil, when used as monotherapy, unfolds antimalarial activity via its metabolite cycloguanil that inhibits dihydrofolate reductase. However, it is proguanil rather than cycloguanil that is responsible for the synergistic interaction with atovaquone (Srivastava, 1999). Proguanil lowers the atovaquone concentration necessary for collapse of the mitochondrial membrane potential (Srivastava, 1999). It was therefore suggested that proguanil inhibits an alternative pathway for generating the mitochondrial membrane potential, which would only become essential once the electron transport chain has been inhibited by atovaquone (Painter, 2007).

Such an alternative pathway could be provided by the F_1_ moiety of the ATP synthase and the ATP^4-^/ADP^3-^ antiporter (Painter, 2007). However, this model has been questioned by the finding that *Saccharomyces cerevisiae* petite (ρ^0^) mutants, which lack complex III and thus rely solely on the alternative pathway to generate their mitochondrial membrane potential, did not show a strong increase in proguanil susceptibility compared to wild type cells (Mounkoro, 2021). Here we investigate the hypothesis that the elusive alternative pathway in *P. falciparum* relies on ATP synthase as a whole, with F_1_ hydrolyzing ATP to export protons through F_O_, thereby generating a mitochondrial membrane potential. *Trypanosoma brucei* bloodstream forms use such a reverse mode of F_O_/F_1_-ATP synthase as their primary way to generate a mitochondrial membrane potential (Nolan, 1992). If complex III inhibition by atovaquone were to force *P. falciparum* to do the same, then we would expect to see synergistic interaction between atovaquone and ATP synthase inhibitors.

Venturicidin A is an antifungal and antimicrobial macrolide from *Streptomyces* spp. that inhibits mitochondrial ATP synthase (Perlin, 1985; McEnery, 1986; Galanis, 1989; Nolan, 1992).

We tested venturicidin A (Santa Cruz sc-202380A), atovaquone (Sigma Aldrich A7986), and proguanil hydrochloride (Sigma Aldrich PHR1713) alone and in combination against *P. falciparum* and *T. brucei*. The 50% inhibitory concentrations (IC_50_) were determined *in vitro* against the asexual stages of *P. falciparum* grown in human erythrocytes using incorporation of [^3^H]hypoxanthine as a readout (Desjardins, 1979), and against axenically grown *T. b. brucei* bloodstream forms with a resazurin-based readout (Räz, 1997). Venturicidin A was active against *P. falciparum* in the submicromolar range (Table 1). Compared to the malaria parasite, the African trypanosomes were hypersensitive to venturicidin A but much less susceptible to atovaquone (Table 1). This is explained by the fact that bloodstream-form *T. brucei* lack complex III (Chaudhuri, 1998; Hellemond, 2005) and solely rely on ATP synthase for the generation of the mitochondrial membrane potential (Nolan, 1992). Based on the obtained IC_50_ values (Table 1), we then tested the combinations proguanil-atovaquone and venturicidin A-atovaquone using a modified form of the fixed ratio isobologram method (Fivelman, 2004; Snyder, 2007). Fractional inhibitory concentrations (FIC) for each combination ratio and each drug were calculated according to (Fivelman, 2004). If for a given combination the sum of the individual FIC (ΣFIC) was below 0.5, it was considered synergistic; a ΣFIC between 0.5 and 2 was considered additive; and combinations with a ΣFIC above 2 were considered antagonistic (Fivelman, 2004).

**Table 1.**
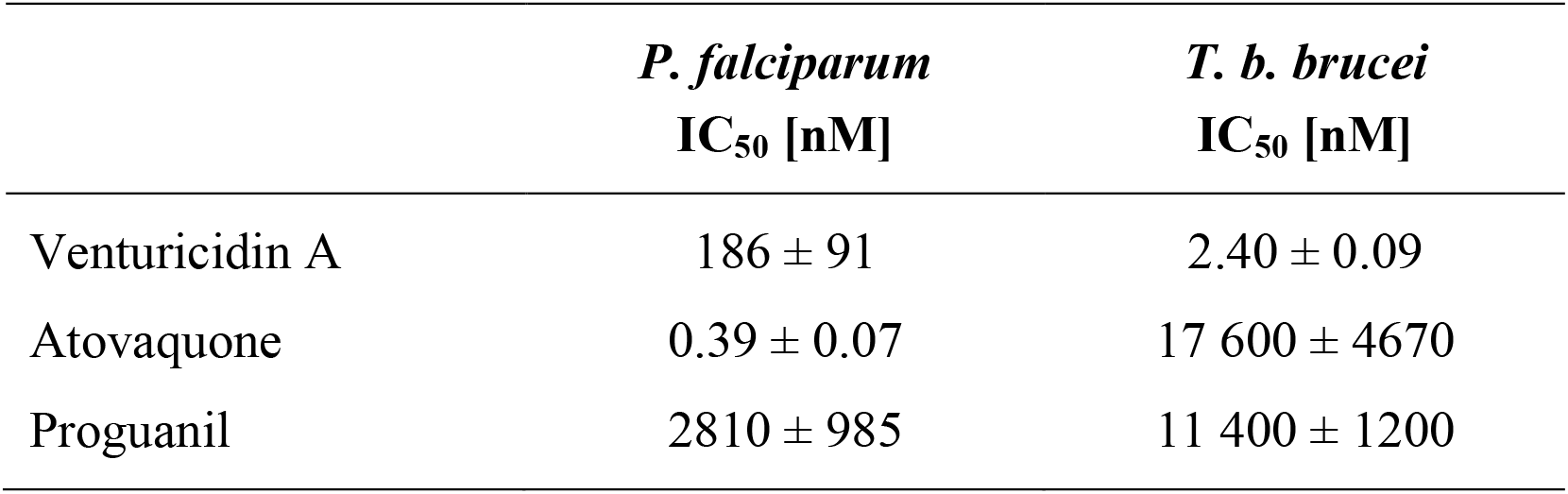
*In vitro* 50% inhibitory concentrations with standard deviation against *P. falciparum* NF54 asexual blood stages and *T. b. brucei* 2T1 bloodstream forms.

|  | *P. falciparum*<br>IC<sub>50</sub> [nM] | *T. b. brucei*<br>IC<sub>50</sub> [nM] |
| --- | --- | --- |
| Venturicidin A | 186 ± 91 | 2.40 ± 0.09 |
| Atovaquone | 0.39 ± 0.07 | 17 600 ± 4670 |
| Proguanil | 2810 ± 985 | 11 400 ± 1200 |

The combination venturicidin A-atovaquone was synergistic against *P. falciparum* (ΣFIC from 0.16 to 0.24; Figure 1A), but not against the *T. b. brucei* bloodstream forms (ΣFIC from 1.08 to 1.42; Figure 1B). We hypothesize that the synergy in *P. falciparum* is the result of simultaneous inhibition of the parasite’s primary and alternative pathways for generating the mitochondrial membrane potential, i.e. complex III inhibition by atovaquone and ATP synthase inhibition by venturicidin A. The absence of a synergistic effect in *T. brucei* is in agreement with this hypothesis, considering that complex III is absent in *T. brucei* bloodstream forms (Chaudhuri, 1998; Hellemond, 2005), and that venturicidin A by itself is able to collapse the mitochondrial membrane potential (Hauser, 2024). Similar results were obtained for proguanil-atovaquone: as expected, the combination was synergistic in *P. falciparum* (ΣFIC from 0.23 to 0.32; Figure 1C) and again, it was not in *T. b. brucei* (ΣFIC from 1.16 to 1.59; Figure 1D). This could indicate a similar mechanism of action as suggested for the venturicidin A-atovaquone combination. It is supported by the finding that proguanil inhibited ATP hydrolysis of the bovine ATP synthase (Bridges, 2014). Interestingly, proguanil only inhibited ATP hydrolysis by the whole F_O_/F_1_-ATP complex, but not by the detached F_1_ moiety (Bridges, 2014), which could explain why the *S. cerevisiae* ρ^0^ mutants, which lack F_O_ as well as complex III, did not show an increased sensitivity to proguanil (Mounkoro, 2021).

**Figure 1.**
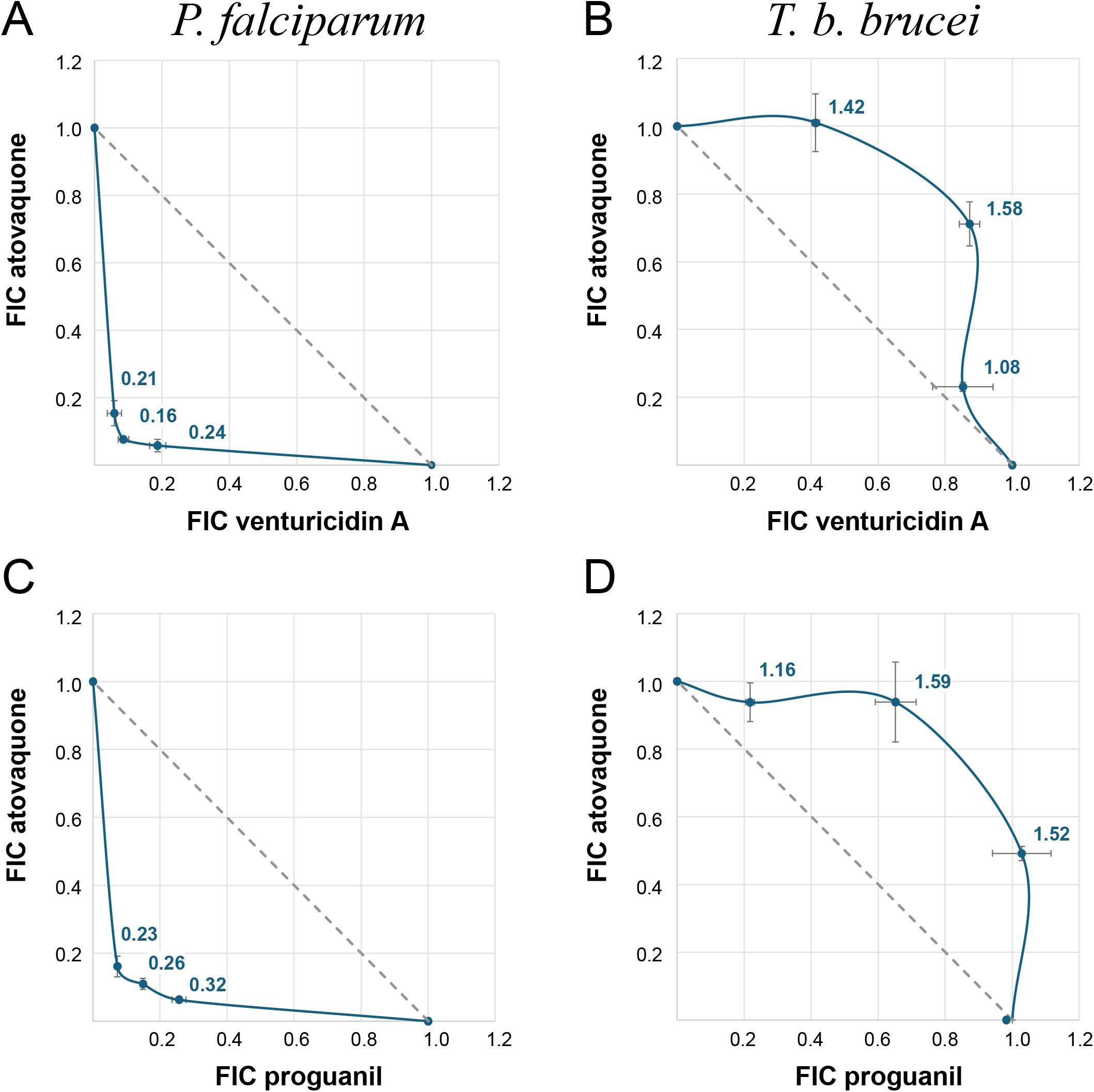
Isobolograms constructed from first combining drugs at fixed ratios of their respective IC_50_ (4:0, 3:1, 2:2, 1:3, 0:4), then determining the IC_50_ for each of these combinations. Fractional inhibitory concentrations (FIC) were calculated according to (Fivelman, 2004). ΣFIC are printed in bold blue, the dashed line depicts ΣFIC=1. The isobolograms with *P. falciparum* clearly indicate synergistic interaction (ΣFIC<0.5). The curves with *T. b. brucei*, in spite of their convex shape, did not meet the threshold (ΣFIC>2) for antagonism. All assays were performed in triplicate; mean FIC are shown with standard deviation.

Our results are consistent with the model that ATP synthase running in reverse mode is the alternative pathway for mitochondrial membrane energization – provided that the malaria parasites actually possess a F_O_/F_1_-ATP synthase! This was not apparent from the *P. falciparum* genome sequence (Gardner, 2002; Vaidya, 2005). However, experimental evidence pointed to a F_O_/F_1_-ATP synthase in the mitochondrion of *P. falciparum* that is functional and essential (Nina, 2011). Subsequently, ATP synthase subunits were identified in *Toxoplasma gondii* (Huet, 2018; Salunke, 2018), many with orthologues in *P. falciparum* (Muhleip, 2021; Maclean, 2022), and the structure of the *T. gondii* ATP synthase showed a clear association between F_O_ and F_1_ (Muhleip, 2021).

In conclusion, we report that venturicidin A is active against intraerythrocytic *P. falciparum*, and that it synergistically interacts with atovaquone. We suggest that the simultaneous inhibition of the primary and the alternative pathway for generating the mitochondrial membrane potential, by complex III and F_O_/F_1_-ATP synthase, respectively, is responsible for this effect. The synergism between proguanil and atovaquone might follow a similar mechanism. These findings further highlight energization of the *P. falciparum* mitochondrion as a target for synergistic antimalarial combination therapy.

## Acknowledgement

We would like to thank the Swiss National Science Foundation for their financial support (grant No. 310030_185163).

